# A novel overlapping gene *azyx-1* affects the translation of zyxin in *C. elegans*

**DOI:** 10.1101/2022.09.09.507294

**Authors:** Bhavesh S. Parmar, Ellen Geens, Elke Vandewyer, Amanda Kieswetter, Christina Ludwig, Liesbet Temmerman

**Affiliations:** Animal Physiology and Neurobiology, University of Leuven (KU Leuven), Belgium; Bavarian Center for Biomolecular Mass Spectrometry (BayBioMS), Technische Universität München, Germany

## Abstract

Overlapping genes are widely prevalent, however, their expression and consequences are poorly understood. Here, we describe and functionally characterize a novel *zyx-1* overlapping gene, *azyx-1*, with distinct regulatory functions in *C. elegans*. We observed conservation of alternative open reading frames overlapping the 5’ region of zyxin family members in several animal species, and find shared sites of *azyx-1* and zyxin proteoform expression in *C. elegans*. In line with a standard ribosome scanning model, our results support *cis* regulation of *zyx-1* long isoform(s) by upstream initiating *azyx-1a*. Moreover, we report on a rare observation of *trans* regulation of *zyx-1* by *azyx-1*, with evidence of increased ZYX-1 upon *azyx-1* overexpression. Our results suggest a dual role for *azyx-1* in influencing *zyx-1* proteoform heterogeneity and highlights its impact on *C. elegans* muscular integrity and locomotion.

## Introduction

Zyxins belong to a subfamily of conserved LIM domain-containing proteins found across eukaryotes and characterized for their role in cell-ECM (extra-cellular matrix) adhesion and cytoskeleton organisation (Beckerle 1997; Zheng and Zhao 2007; Sadler et al. 1992; Freyd et al. 1990). Characterized by a proline-rich N-terminus and three consecutive LIM domains in their C-terminal region (Sadler et al. 1992; Beckerle 1997; Kadrmas and Beckerle 2004), zyxins regulate actin assembly and remodelling, as well as cell motility (Fraley et al. 2012; Hoffman et al. 2006; Nix and Beckerle 1997; Winkelman et al. 2020). In line with this, human zyxin is implicated in stretch-induced gene expression changes via active nuclear translocation (Wójtowicz et al. 2010). Moreover, it also promotes apoptosis in response to DNA damage (Crone et al. 2011).

While multiple distinct zyxin proteins are present in vertebrates (for example Lpp, Trip6 and Zyx), *C. elegans* contains an unique zyxin gene, *zyx-1*, with five annotated protein isoforms, of which isoforms a and b are predominantly expressed (Kadrmas and Beckerle 2004; Lecroisey et al. 2013; Luo et al. 2014). There are anatomical differences in isoform expression, with ZYX-1b observed in body wall muscle, pharynx, vulva, spermathecae and multiple neurons, whereas ZYX-1a mainly localises to tail phasmid neurons and uterine muscle, and weakly so, body wall muscle (Luo et al. 2014). In *C. elegans, zyx-1* has been postulated to have a minor role in reproduction, however, the mechanism(s) and isoform(s) involved remain elusive (Castaneda et al. 2021; Smith et al. 2002). Beyond this, *C. elegans zyx-1* is hypothesized to be functionally analogous to vertebrate zyxin, with LIM domains acting as mechanical stabilizers at focal adhesions, and the proline-rich N-terminus involved in sensing muscle cell damage (Lecroisey et al. 2013).

Previous studies revealed that only the ZYX-1b isoform regulates synapse maintenance and development, while in the context of a dystrophic mutant background, the longer ZYX-1a isoform partially rescues muscle degeneration in an ATN-1 dependent manner, highlighting how not only expression patterns, but also molecular functions are isoform-dependent for *C. elegans* zyxin (Luo et al. 2014; Lecroisey et al. 2013). At a gene-regulatory level, this use of alternative splicing products and functional diversification of proteoforms for *C. elegans* zyxin is in line with observations made for several LIM domain proteins across eukaryotes (Zheng et al. 2016).

In general, alternative and overlapping open reading frames (ORFs) arising out of polycistronic mRNA can contribute to post-transcriptional regulation (Brunet et al. 2020; Makalowska et al. 2005; Wright et al. 2021). A recent community-wide effort for annotation of such genomic loci categorized overlapping genes based on their initiation and termination codon with respect to the main coding sequence of a given transcript (Mudge et al. 2021). Of these, ORFs with an upstream start site (upstream or upstream overlapping; uORF and uoORF) often influence the translation of the main coding sequence, based on the evidence of the ones that have been detected and investigated in detail (Chew et al. 2016; Schulz et al. 2018; Zhang et al. 2019; Young and Wek 2016; Calvo et al. 2009). From a more human-centred future perspective, uORFs are a rather unexplored niche for translational research: with a predicted prevalence in over 50% of human genes and first examples regulating translation of disease-associated genes already emerging (Lee et al. 2021; Schulz et al. 2018), the field is bound to not only lead to more fundamental, but also application-oriented insights. Keeping this broader context in mind, we here focus on more fundamental principles of uORFs in a model organism context.

We previously provided mass spectrometric evidence for 467 splice variants and 85 non-canonical gene products, including from polycistronic and ncRNA translation, in *C. elegans* (Parmar et al. 2021). Of these, one newly discovered gene, *azyx-1* (<u>a</u>lternative N-terminal ORF of *<u>zyx</u>-1*), was identified as an 166 amino acid-long protein, translated from the 5’UTR of *zyx-1*. In this study, we provide evidence for two protein isoforms of *azyx-1*, one initiating upstream and another downstream of *zyx-1* AUG, and both overlapping the proline-rich N-terminus of ZYX-1 long isoforms. To understand whether zyxin proteins could also be regulated by upstream/overlapping ORFs, we here explore functional relevance of *azyx-1* and its relation to zyxin in the model organism *C. elegans*.

## Results

### A novel gene, azyx-1, reveals putative syntenic conservation of overlapping genes on eukaryotic zyxin

The *C. elegans* genome contains 2468 predicted ORFs initiating in 5’ UTRs, however, only 8 of those are supported by mass spectrometric evidence (Parmar et al. 2021; Brunet et al. 2018). One of these is *azyx-1*, a non-canonical gene overlapping the gene encoding the only *C. elegans* zyxin, *zyx-1* (Parmar et al. 2021). *azyx-1* is translated from a different reading frame than *zyx-1*, starting from an AUG 184 bp upstream of the *zyx-1* initiation site and delivering a polypeptide product with a sequence length of 166 amino acids (AZYX-1a, Fig. 1A). The same locus also embeds a putative shorter AZYX-1 isoform (AZYX-1b), initiating within the *zyx-1* coding sequence and containing only the 106 C-terminal amino acids of AZYX-1a (Fig. 1A). Six different transcripts have been described for the *zyx-1* locus (F24G4.3, WormBase version WS283): two are associated with isoform a, and then a single transcript for isoforms b-e each (Fig. 1A). Based on these, AZYX-1a can be translated from the F42G4.3a.1 transcript which contains a longer 5’UTR, whereas AZYX-1b could be translated from all three transcripts of the longer *zyxin* isoforms a and e: F42G4.3a.1, F42G4.3a.2 and F42G4.3e.1. We previously mapped unique peptides to *azyx-1*, covering >60% of the 166 amino acids-long protein sequence at 1% FDR (Parmar et al. 2021), showing that *azyx-1* indeed is translated. Since LIM-domain proteins, to which zyxins belong, often result from alternative splicing and display strong domain conservation across eukaryotes (Zheng et al. 2016), we asked whether *azyx-1* or the phenomenon of overlapping genes at zyxin loci is also conserved across eukaryotes. To that end, we searched for predicted ORFs initiating in 5’ UTRs (uORF and uoORF) or within the coding sequence (oORF) of zyxin orthologs (*lpp, trip6* and *zyx*) in 8 animals (OpenProt release 1.6, Supplemental Table. S4). We identified 14 ORFs initiating upstream of *zyx-1* orthologs in 7 species including *C. elegans* (Supplemental Table. S4, Fig. 1B). Moreover, 6 of these 5’ UTR initiating ORFs were conserved in vertebrates as alternative oORFs within the coding region of zyxin orthologs (Supplemental Table. S4). Together, our observations suggest a syntenic conservation of overlapping genes towards the 5’ end of zyxin orthologs, in combination with the widely prevalent alternative splicing observed in zyxins.

**Figure 1.**
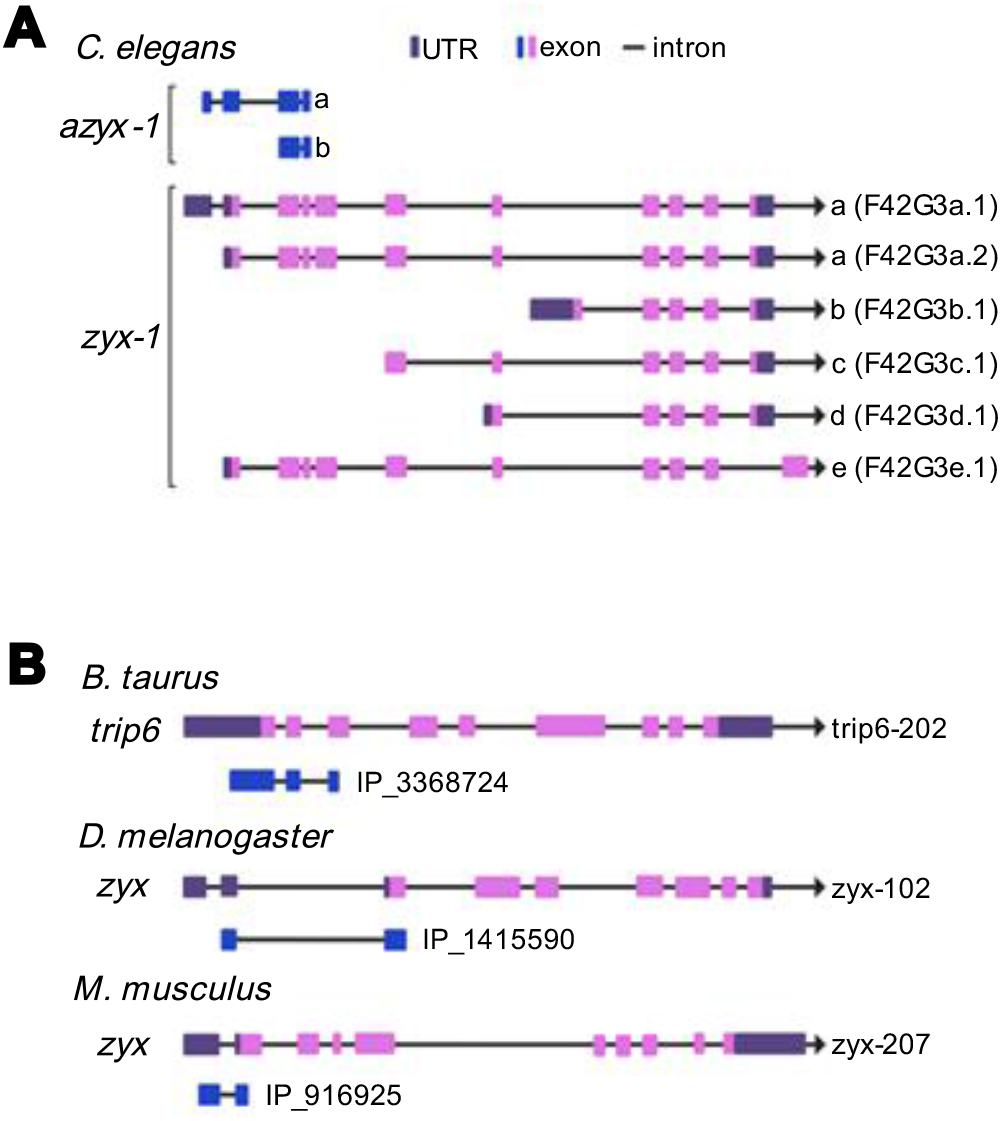
Schematic representation of zyxin loci including overlapping uoORFs. Exons (pink: canonical, blue: uoORF/oORF), introns (line) and UTRs (purple) indicated. All loci are shown according to the direction of transcription (arrowhead). **A**. Six zyx-1 transcripts are recognized in C. elegans, corresponding to five isoforms. The coding sequence for the uoORF/oORF (blue) azyx-1 as identified previously by Parmar et al. 2021 is contained in the long transcripts. **B**. Examples of uoORFs (blue) predicted by OpenProt (Brunet et al. 2018) with respect to zyxin orthologs of B. taurus, D. melanogaster and M. musculus. The gene names (left), transcript ID (right) and OpenProt IDs are provided next to each schematic representation.

### *azyx-1* and *zyx-1a* reporters are observed in partially overlapping anatomical locations

To understand which cells or tissues of *C. elegans* may express *azyx-1*, we generated an extrachromosomal *azyx-1* reporter strain (LSC1959). While the *zyx-1*a start codon is contained within this sequence, it doesn’t share the *azyx-1* reading frame. Hence, the *azyx-1::mNeonGreen* fusion construct cannot lead to a fluorescent signal should translation initiate at the downstream *zyx-1*a start codon. We observed a strong fluorescent signal in body wall muscle, vulval muscle and very faintly in unidentified structures in the head and tail (Fig. 2A-D). This fluorescence pattern was consistent, as observed in L4 and adult worms (Supplemental Fig. S1A-D, Fig. 2). Next, we looked for expression of *zyx-1* isoforms as reported via extrachromosomal arrays by Lou *et al*. (2014). In line with their report, we also observe the *zyx-1* long isoform predominantly in tail neurons, and faintly in body wall muscle, and additionally also faintly in the pharynx, and more brightly in an unidentified structure in the head (Fig. 4A, middle panel of control). Interestingly, the short *zyx-1b* isoform is also strongly expressed in body wall and vulval muscle (Luo et al. 2014), where we observe *azyx-1* expression (Fig. 2A). Thus, beyond alternative splicing in zyxins, our observations suggest that in those locations, proteoform diversification and localisation fine-tuning may putatively also occur via polycistronic transcripts, with the observed expression patterns suggesting the possibility of mutually exclusive translation from long *zyx* transcripts.

**Figure 2.**
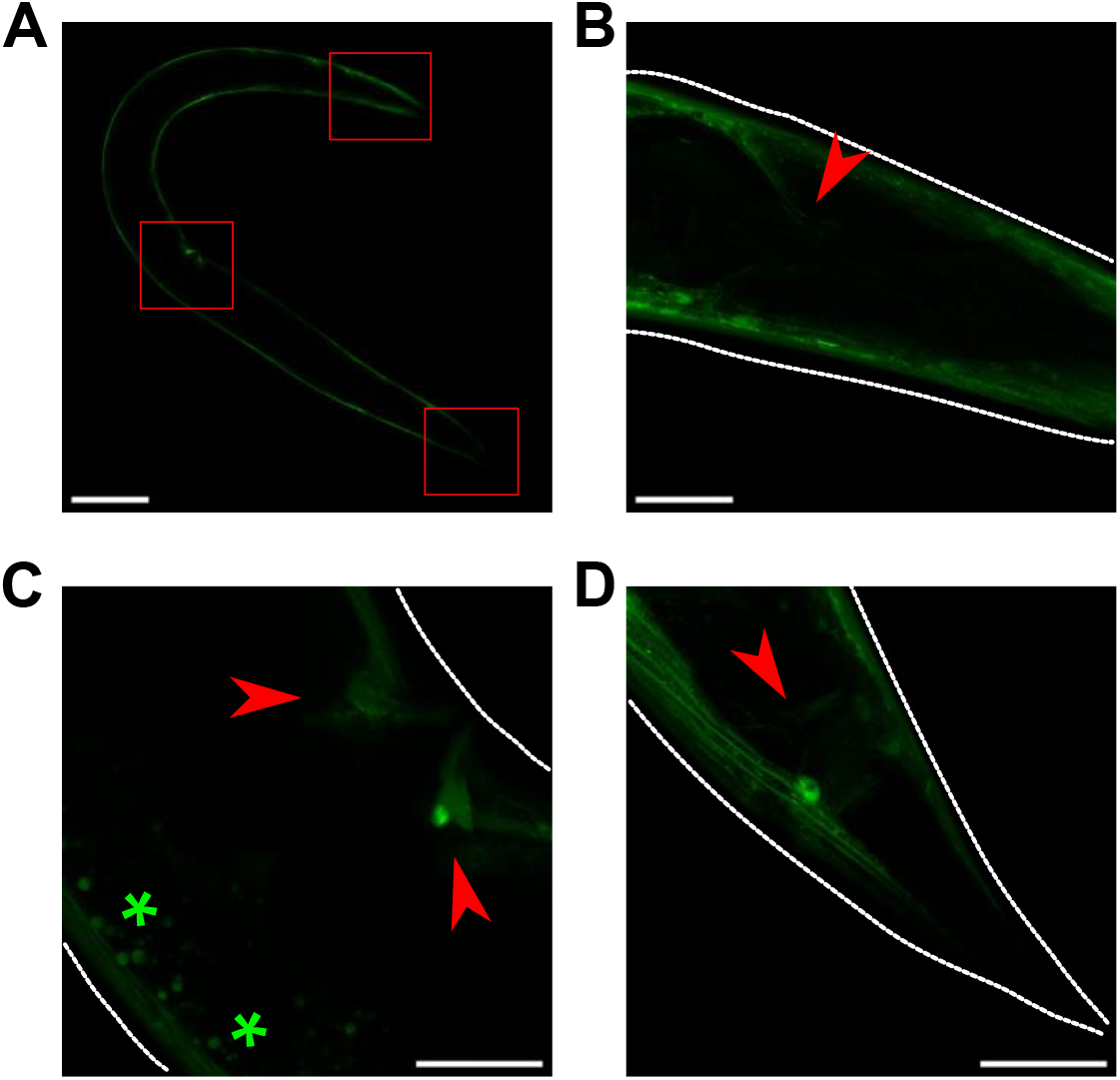
azyx-1 is prominently expressed in body wall and vulval muscle and faintly in the head and tail region. Anterior always to the left. **A-D**. For azyx-1 localisation, azyx-1p::azyx-1::mNeonGreen:: azyx-1 3’UTR was expressed extra-chromosomally in a wild type background. The reporter protein is clearly visible in **(A)** body-wall muscle (scale bar - 100 μm), with red boxes indicating **(B)** two unidentified neurites in the head, **(C)** vulval muscle, and **(D)** in the tail region (scale bar - 20 μm, autofluorescence *). For L4 life stage, see Supplemental Fig. S1A-C.

### AZYX-1 and ZYX-1 levels vary with age

To understand which protein products are generated by these overlapping genes, we quantified AZYX-1 and ZYX-1 at three different ages: L4 larvae, and young (day-1) and post-reproductive (day-8) adults. We selected these life stages based on *azyx-1* and *zyx-1* expression in body wall muscle (Fig. 2), combined with the knowledge that *C. elegans* musculature deteriorates with age (Herndon et al. 2002). We observed a relative increase of about 4-fold for AZYX-1 (*p = 8*.*3e-5*) and 6-fold for ZYX-1 (*p = 1*.*9e-5)* in day 1 adult *vs* L4 stages (Fig. 3A, Supplemental Fig. S2). While AZYX-1 at day 8 of adulthood remained high (*p = 0*.*77*, Fig. 3A, Supplemental Fig. S2A), ZYX-1 of these post-reproductive animals was intermediary between L4 and day 1 adult levels (Fig. 3A). By relying on quantitative data of the specific peptides making up these proteins, we can further deduce that changes in adult zyxin levels differ for different proteoforms: at day 8 *vs* day 1 of adulthood, 2 out of 3 quantified N-terminal peptides (ZYX-1.1 and 1.3) were comparable to day 1 levels, while the quantified C-terminal peptides (ZYX-1.4 and ZYX-1.5) significantly reduced (*p = <0*.*02*, Fig. 3B, Supplemental Fig. S2B). For AZYX-1, 5 out of 7 peptides remain unchanged while 2 (shared by both AZYX-1a and AZYX-1b) show a decline (Fig. 3C). Together, these data suggest that AZYX-1 levels remain stable while ZYX-1 levels decline between day 1 and day 8 of adulthood. This is corroborated by similar transcript level fold change of *zyx-1* for day 1 and day 8 adulthood (Roux et al. 2022). Furthermore, it appears that shorter isoforms of *zyx-1* may reduce with age, whereas the longer isoforms might be less susceptible to such a post-reproductive decline.

**Figure 3.**
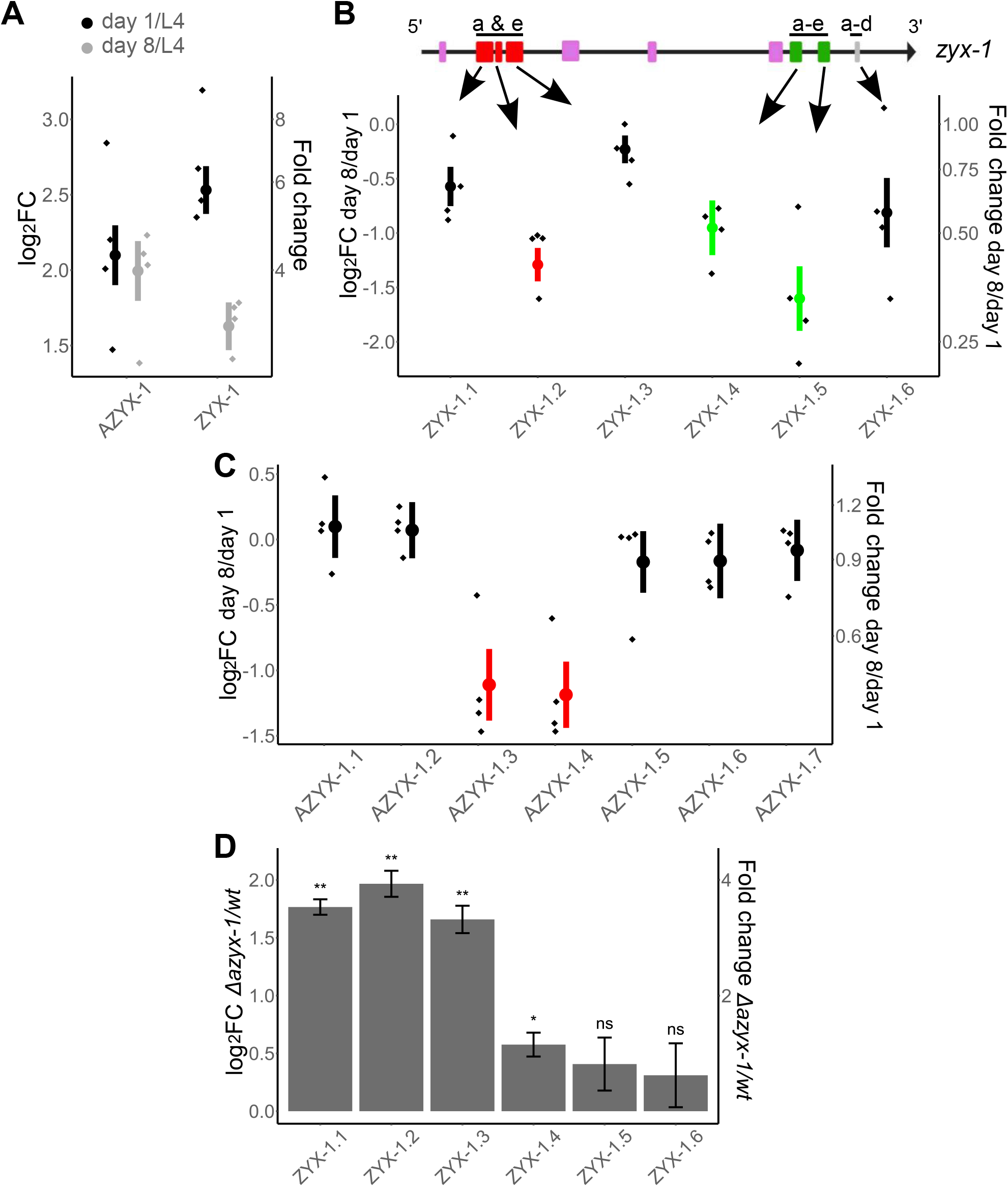
Targeted quantitation in aging worms and azyx-1a mutant reveals age dependent zyx-1 fold change and cis regulation by azyx-1. Graph axes indicate log2 fold change (left) and absolute fold change (right). Data normalized to a spike-in peptide (1 fmol/worm) for aging wt worms and GPD-3 for mutant vs wt comparison, with 4 biological replicates (diamonds) for each time point, mean (circle) and standard error bars. **A**. Fold change at day 1 (black) and day 8 (grey) of adulthood in comparison to L4 larval stage for all quantified peptides of AZYX-1 (7 peptides) and ZYX-1 (6 peptides) combined. **B**. Fold change at day 8 in comparison to day 1 of adulthood for individual ZYX-1 peptides (1.1 to 1.6). zyx-1 gene model indicates the location of peptides unique to long isoform a & e (1.1-1.3) and peptides shared by longer and shorter isoforms (1.4-1.6). **C**. Fold change at day 8 in comparison to day 1 of adulthood for individual AZYX-1 peptides (1.1 to 1.7). AZYX-1.1 and 1.2 are unique to AZYX-1a while the rest are shared by AZYX-1a and AZYX-1b. Significantly downregulated peptides are coloured as per their position in the gene model (Red: long, green: shared; fold change cut-off = <0.66, p-value = <0.02). **D**. Peptides unique to ZYX-1 long isoforms (ZYX-1.1-1.3) show a consistent increase in the mutant, while peptides shared between all ZYX-1 (1.4-1.6) isoforms remain similar to wildtype levels (at day 1 of adulthood). p-value: * <0.05, ** <0.01, *** <0.001, ns = not significant, n = 4 biological replicates, with data normalized to GPD-3. For spike-in and HIS-24 normalization see Supplemental Fig. S3A and B.

### *azyx-1* likely exhibits *cis* control over *zyx-1*

Since *azyx-1* initiation lies 184bp upstream of *zyx-1*, we hypothesized a *cis* regulation of downstream *zyx-1* major ORF by *azyx-1*. To test whether this could be the case, we generated a *azyx-1a* mutant (LSC1898) and quantified ZYX-1 peptides. We observed a substantial increase in N-terminal ZYX-1 peptides (ZYX-1.1-1.3) for *azyx-1* mutants *vs* wild type (Fig. 3D). Peptides that were not specific to the long zyxin proteoforms (ZYX-1.4-1.6) showed a much more modest change – or even none at all – in mutant *vs* control. The N-terminal peptides, specific to *zyx-1* long isoforms (ZYX-1a/e), showed a 2-3-fold range increase upon *azyx-1a* mutation. This was observed for three independent means of data normalization (GDP-3, spike-in, HIS-24; Fig. 3D Supplemental Fig. S3A-C), supporting the solidity of the claim that removal of the *azyx-1a* start codon increases zyxin levels in a proteoform-biased way. Interestingly, AZYX-1a start codon removal led to suppression of all downstream peptides shared by AZYX-1b, except one, which was significantly down-regulated. This drop below the limit of detection of AZYX-1a and b shared peptides suggests that AZYX-1a might be the prominently translated isoform (Supplemental Fig. S3D). Together, our results fit the hypothesis that *azyx-1* may inhibit translation of downstream *zyx-1* isoforms, likely affecting the longer isoforms more due to a shared transcript. This does not rule out the possibility of trans regulation (see below) or of an internal ribosome entry site (IRES) downstream of the AZYX-1a start codon for ZYX-1a initiation, however, based on *in silico* analysis, there is no evidence of IRES within the region downstream of the AZYX-1a start site.

### *azyx-1* overexpression increases ZYX-1 levels and accumulation in GABAergic motor neurons

While we suspect that uoORF *azyx-1a* may exhibit *cis* control over translation of *zyx-1* long isoforms, measurable quantities of AZYX-1 peptides are being produced *in vivo* (Fig. 2 and 3, Parmar et al. 2021). Given the detection of two isoform-specific peptides (Supplemental Table. S2), these certainly are translation products of *azyx-1a*. A putative shorter isoform, *azyx-1b*, with a predicted start codon downstream of that of the *zyx-1* long isoforms could also contribute to the remainder of the measured signal (Fig. 1A, 2 and 3, Parmar et al. 2021). Therefore, we asked whether *azyx-1* might also influence *C. elegans* zyxin in *trans*. We made use of a reporter system that fuses all zyxin isoforms to GFP, and only the long ones (a/e) additionally to mCherry (Luo et al. 2014) and extrachromosomally expressed *azyx-1* (Fig. 4A). We measured a significant increase in all (*zyx-1::GFP*), as well as in the long (*mCherry::zyx-1*) proteoform reporter levels specifically, upon *azyx-1* overexpression in the reporter strain (*p<0*.*001*; Fig. 4B-C). This observation suggests that production of ZYX-1 may be stimulated by the presence of AZYX-1. In addition, the ratios for mCherry/GFP show no significant differences, suggesting an overall ZYX-1 increase under forced overexpression of *azyx-1*, irrespective of proteoform (Fig. 4D). So far, our results support the presence of both *cis* and *trans* control of *zyx-1* by *azyx-1 in vivo*.

**Figure 4.**
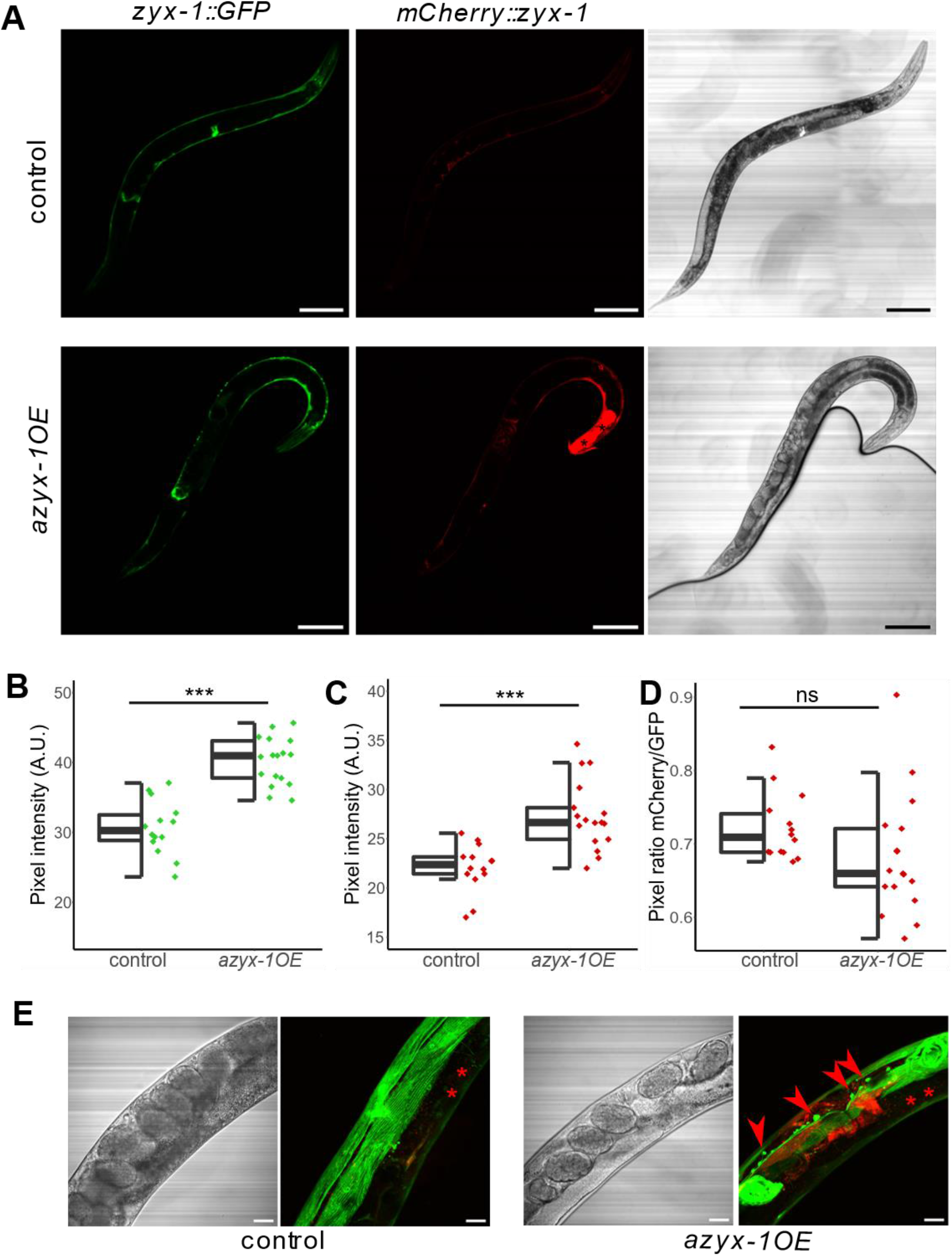
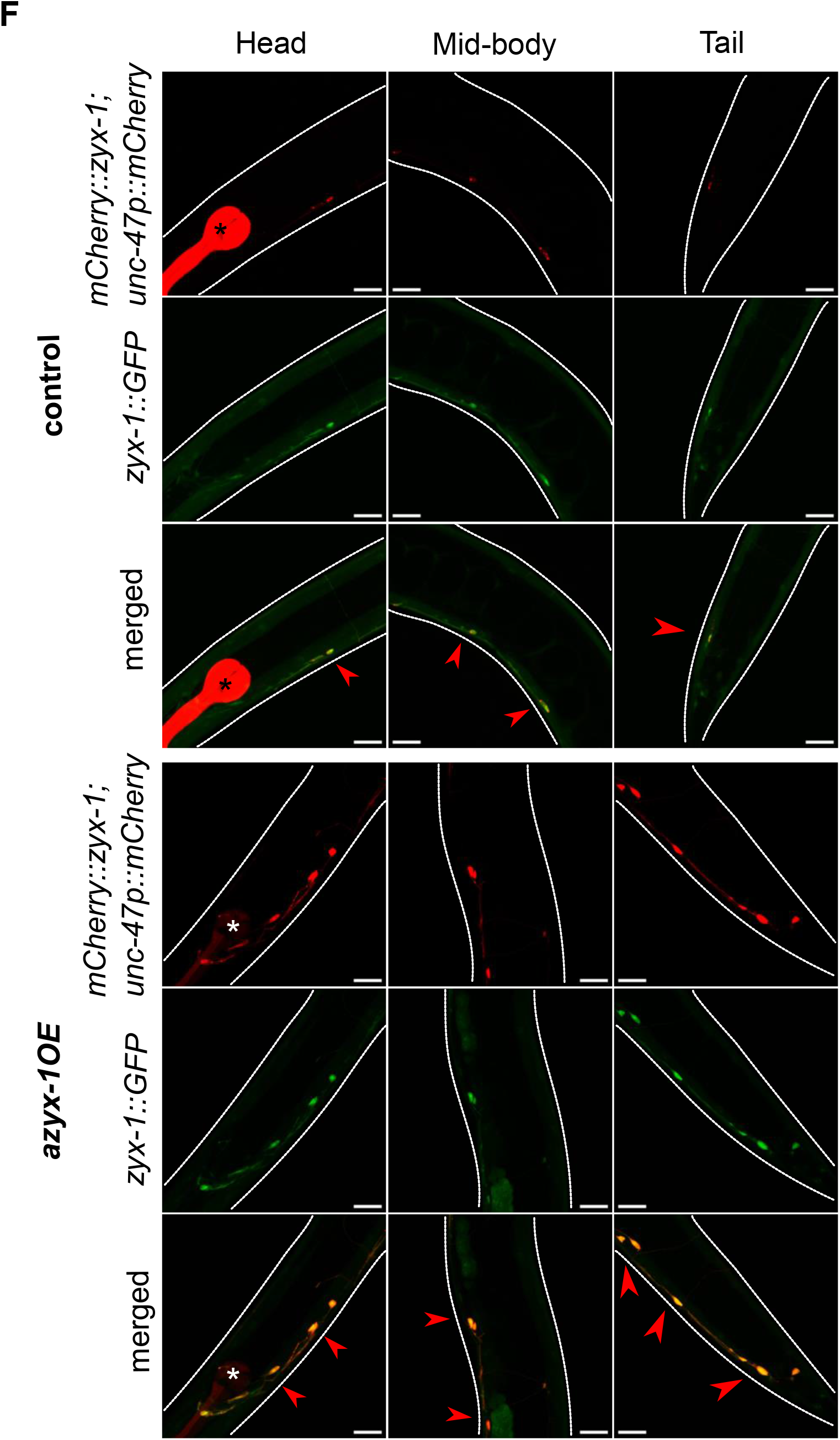
azyx-1 overexpression increases zyxin reporter signal and leads to zyxin accumulation in motor neurons. The integrated reporter control (LSC1870) **A. (upper**) contains ZYX-1::GFP (representing all isoforms) and mCherry::ZYX-1 (representing long isoforms), which is compared against a strain with the same genetic reporter background **A. (lower**), extrachromosomally overexpressing azyx-1 (azyx-1OE, LSC1960). Anterior to the right, scale bar 100 μm. **B**. GFP (p=4.85e-8) and **C**. mCherry (p=4.82e-5) quantification showing increase in signal upon azyx-1 overexpression, with * indicating significance (ANOVA) for n≥14 per condition. **D**. ratio of mCherry/GFP. **E**. increased accumulation of GFP along the ventral nerve cord as observed in azyx-1OE (LSC1960) compared to control reporter (LSC1870), *intestinal autofluorescence and **F**. Co-localization (merged, red arrows) of zyx-1::GFP in motor neurons upon azyx-1 overexpression as imaged in adult head, mid-body and tail. Upper panels: control worms only expressing the neuronal reporter construct (unc-47p::mCherry) and *co-injection marker (myo-2p::mCherry), lower panels: azyx-1 OE in the control background. Scale bars: 20 μm.

Upon *azyx-1* overexpression, we also observed an increase in GFP signal aggregation along the ventral nerve cord in what appeared to be motor neurons (Fig. 4E). Colocalization with *unc-47p::mCherry*-positive cells identifies GABAergic neurons along the ventral nerve cord as sites of this *zyx-1* accumulation under *azyx-1* overexpression (Fig. 4F). *zyx-1* has previously been reported to be expressed in GABAergic neurons (Spencer et al. 2011), and its accumulation in these motor neurons upon increased AZYX-1 levels made us hypothesize that *azyx-1* overexpressors may suffer locomotion-related impairments.

### *azyx-1 – zyx-1* affect muscle integrity and neuromuscular behaviour in *C. elegans*

Our expression analysis revealed that *azyx-1* and *zyx-1* are abundantly expressed in body wall muscle (Fig. 2A and E). In line with this, using blinded and randomized manual scoring (Fig. 5A), we observed a quantifiable loss of muscle fibre integrity in *azyx-1* overexpression conditions, as compared to control animals (Fig. 5B). To test whether manipulating AZYX-1 levels could, as is already known for *zyxin* deletion (Lecroisey et al. 2013), affect muscle performance, we performed burrowing assays (Lesanpezeshki et al. 2019). Here, the genetic removal or addition of *azyx-1* both resulted in defective burrowing as compared to wild type (two-way ANOVA *p=0*.*0025*, Tukey HSD *p=0*.*01 and 0*.*047 respectively*), and the deletion mutant phenotype could be rescued by extrachromosomal resupplementation of *azyx-1*, whereas no significant difference was observed between *azyx-1* conditions and the positive control *zyx-1(gk190)* (Fig. 5C).

**Figure 5.**
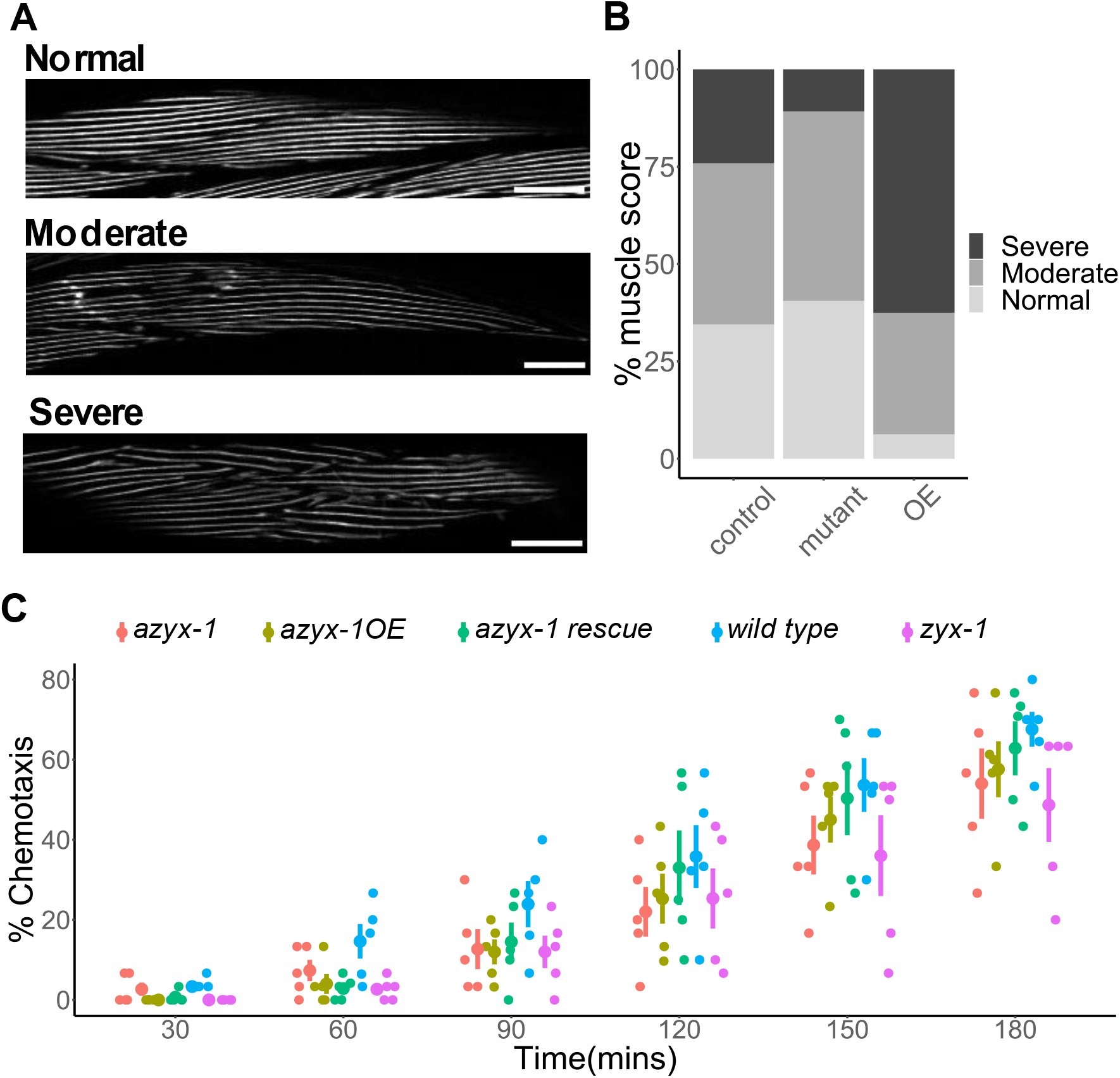
azyx-1 affects muscle integrity and burrowing behaviour. A fluorescent myo-3 reporter (RW1596) was used and either crossed with an azyx-1 deletion mutant (LSC2001) or injected to create an overexpressor (LSC2000) to score muscular quality via blinded image analysis (n≥ 25 per condition). **A**. Example images of what was scored as normal, vs a moderate or severe muscular defect. **B**. Distribution of observed muscular phenotypes in azyx-1 mutant and overexpressor (OE) as compared to controls. **C**. Chemotaxis index of burrowing assay, as cumulatively observed over 180 mins for azyx-1 mutant, rescue and overexpressor in comparison to positive (zyx-1) and negative (wt) controls, n=30 per condition in 5 replicates (30 × 5); p-value two-way ANOVA <0.0025 for strain and time. TukeyHSD wt vs azyx-1OE/azyx-1 mutant/zyx-1 = 0.047,0.01,0.002 respectively.

## Discussion

Although non-canonical ORFs are widely prevalent in eukaryotic genomes (Chen et al. 2020; Ingolia et al. 2014; Bazzini et al. 2014), they often tend to be species-specific (Makalowska et al. 2005; Veeramachaneni et al. 2004; Soldà et al. 2008). These ghost genes often are not susceptible to BLAST analysis due to their small size and inherent genomic variation over species. Upon manually examining the genomic loci of zyxin orthologs, we found evidence of putative syntenic conservation of *azyx-1* across 7 species (Fig. 2B, Supplemental Table. S4). This exemplifies how non-canonical ORFs can escape functional annotation, as the existing automated means for orthology mapping are restricted to (large) sequence similarity. Naturally, overlapping genes are more likely to be conserved across species, as a consequence of parent gene conservation. It is interesting however, that overlapping genes could also be harboured within specific gene isoforms to potentially regulate gene expression at a proteoform level, which we propose is the case for *azyx-1a* and *zyx-1a*.

Since *azyx-1a* and *zyx-1a* likely share a transcript, we asked whether there might be a preference for translation initiation from either of the two start codons. Upon *azyx-1a* start-codon mutation, we observed a substantial increase in ZYX-1 long isoform abundance, while the downstream short isoforms largely remain unaffected (Fig. 3D, Supplemental Fig. S3A-B). Given the global linearity of eukaryotic ribosomal scanning and translation (Vassilenko et al. 2011; Hinnebusch 2011), our results suggest that *azyx-1a* exhibits *cis* control over *zyx-1* long isoform(s), by occupying the scanning ribosome in +2 reading frame. While we cannot fully exclude a direct effect on *zyx-1a* transcription due to this 27bp deletion, our observations are in coherence with the widespread *cis* repression of downstream major ORF translation by upstream ORFs, predicted across vertebrates and observed in cells and animal models (Ribone et al. 2017; Vattem and Wek 2004; Johnstone et al. 2016).

Interestingly, *azyx-1* also contains a downstream oORF isoform, prompting a putative *trans* function. One option would be for AZYX-1 to drag the remaining transcript to nonsense mediated decay (NMD), as has been observed for other uoORFs in plants, yeast and mammalian cells (Nyikó et al. 2009; Hurt et al. 2013; Mendell et al. 2004; Gaba et al. 2005; Johansson and Jacobson 2010). However, *zyx-1* is not present in the available *C. elegans* resource for NMD targets (Muir et al. 2018) and contrary to such expectations, we observe a positive *trans* effect of *azyx-1* on *zyx-1* (Fig. 4). We therefore propose there may be a feed-forward loop that regulates *zyx-1* (long and short) translation via *azyx-1* (Fig. 6). This is in any case a rare observation of reinforcing *trans* control by an upstream (overlapping) gene, which based on our observations is opposed to its proposed uoORF-mediated *cis* effect (Fig. 3D). A similar observation of a combined *cis* and *trans* regulation has been made before for human hepatitis B virus (Chen et al. 2005), and it is possible that the regulatory context could be proteoform-specific. While the *cis* regulation can be explained by standard ribosomal scanning, the molecular interplay involved in sensing AZYX-1 to then regulate *zyx-1* in *trans* remains to be investigated. Since *azyx-1* extends 2475bp into the *zyx-1 long* ORF, it is unlikely for ribosomes to re-initiate at *zyx-1 long* start codons after being released from the *azyx-1* termination site. We hypothesize that the *zyx-1-*overlapping C-terminus of *azyx-1* contains a functional *trans* domain responsible for the feed-forward loop. Previous studies have shown physical interaction between overlapping genes (Bergeron et al. 2013; Klemke et al. 2001), but the mechanisms by which this occurs often remain undiscovered. Our results suggest that *azyx-1/zyx-1* is an interesting candidate to elucidate such interactions and intragenic *trans* regulation.

**Figure 6.**
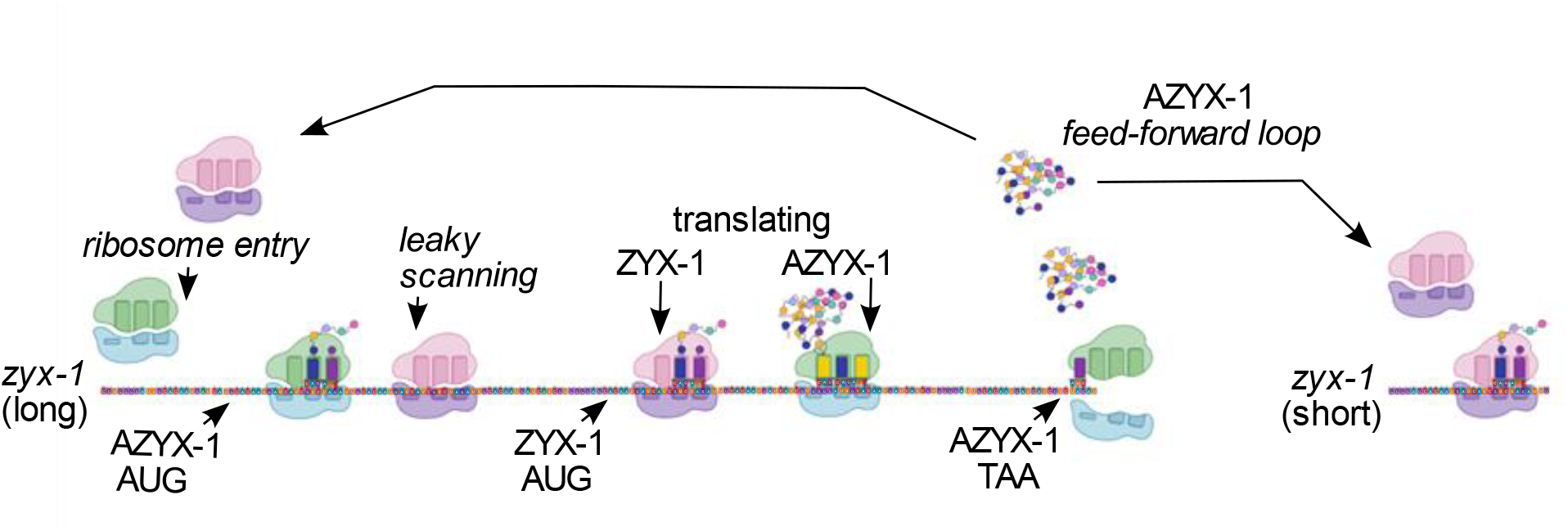
Proposed model of cis and trans regulation of zyx-1 by azyx-1. Ribosomes upon entry onto zyx-1 (long) mRNA initiate translation (green) at uoORF AZYX-1a AUG in +2 reading frame, thus repressing translation of downstream ZYX-1 in cis. Leaky scanning through upstream AZYX-1a initiation leads to ZYX-1 translation (pink). Accumulation of translated AZYX-1 causes increased translation of zyx-1 (long and short) mRNA in a feed-forward loop via an unknown signalling cascade.

Phenotypically, decreased burrowing efficiency in *azyx-1* mutant and overexpressor strains further corroborates our hypothesis of feed-forward loop between *azyx-1* and *zyx-1* (Fig. 5). Moreover, increased localisation of *zyx-1* in motor neurons (Fig. 4G), defective muscle spindles and burrowing chemotaxis upon *azyx-1* overexpression (Fig. 5) suggest *azyx-1* as a key player in *zyx-1*-mediated muscular integrity and locomotion. Since *zyx-1* shorter isoforms (b and d) are mainly expressed in body wall muscle and motor neurons (Luo et al. 2014; Lecroisey et al. 2013), we anticipate that the observed increase upon *azyx-1* overexpression corresponds to shorter and not *zyx-1* long isoforms (Fig. 5B).

Previous studies observed substantial decrease in worm muscle degeneration upon RNAi knockdown of 5’ region of *zyx-1a* and 3’ of *zyx-1* in dystrophic background (*dyc-1;hlh-1*), with a distinct isoform involvement (Lecroisey et al. 2013). Our results show that *azyx-1* overexpression leads to *zyx-1* increase and muscle deterioration, while a *azyx-1* mutant has significantly increased *zyx-*1 long isoforms and does not suffer measurable muscular defects (Fig. 5B). We hypothesize that the observed effect stems from two distinct pathways, which may or may not involve direct modulation by *azyx-1*. The *zyx-1 long* isoform, with its characteristic N-terminal proline rich region, has been hypothesized to translocate to the nucleus under cytoplasmic stress and activate transcription of genes involved in repair of damage (Lecroisey et al. 2013). This would explain the muscular integrity of the *azyx-1* mutant, possibly driven by its *zyx-1* long isoform increase. On the other hand, *azyx-1* overexpression could be causing an imbalance in long/short *zyx-1* ratio via excess LIM domain expression and consequent disturbance of endogenous *zyx-1* homeostasis, similar to that observed previously upon LIM domain overexpression in Vero cells (Nix et al. 2001). Conversely, *azyx-1* could harbour a hitherto unknown functional domain, which upon overexpression, causes loss of muscular integrity. Given the conservation of LIM domain proteins across eukaryotes, and the evidence of putative syntenic conservation for *azyx-1* (Fig. 1), we believe this could be an interesting avenue for future research on LIM domain proteins and prevalence of similar overlapping genes in eukaryotic systems.

## Materials and methods

### Reagents and tools table

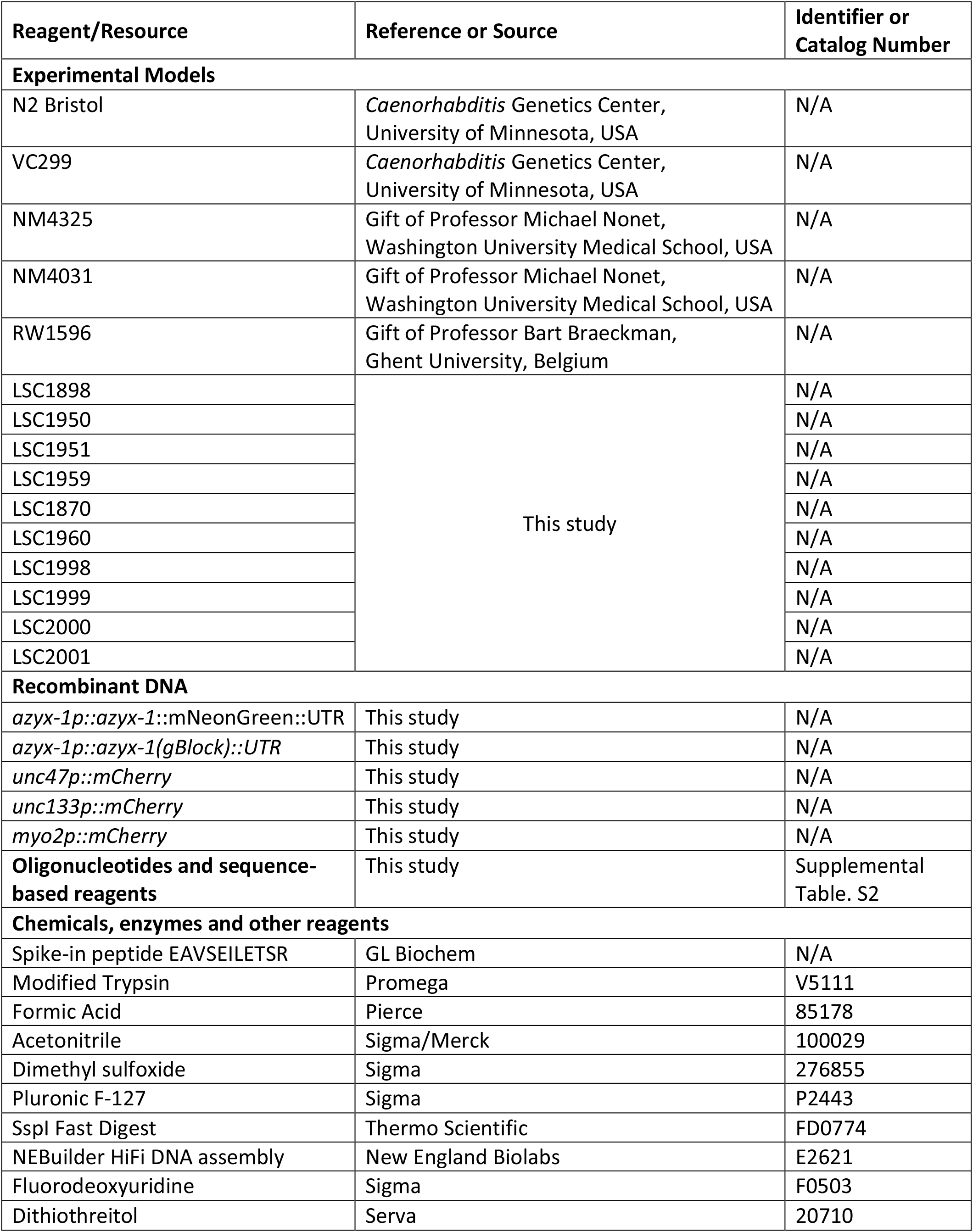
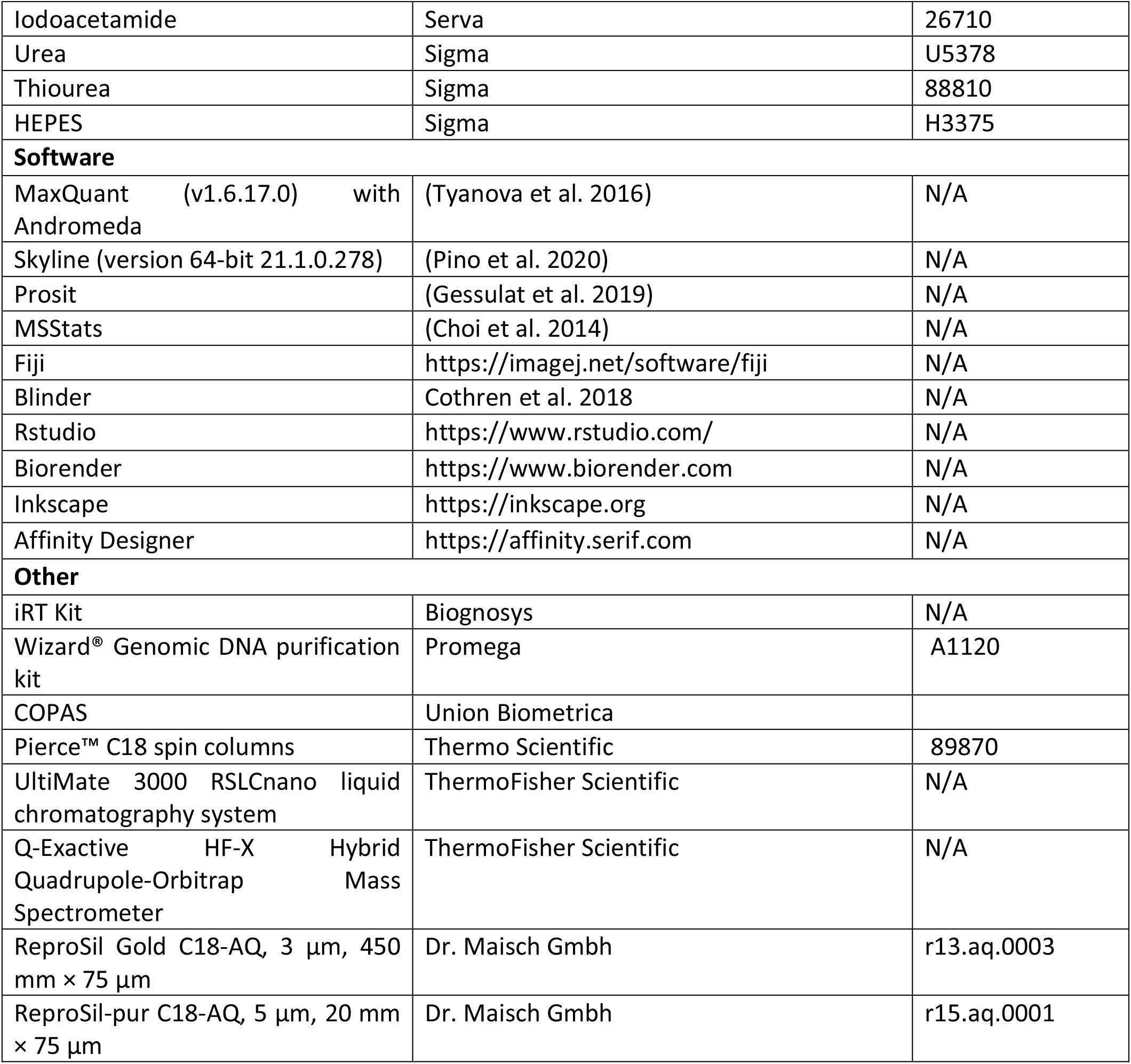

### Worm culture

All strains used in this study (Supplemental Table. S1) were cultured at 20 °C on nematode growth medium (NGM) plates seeded with *E. coli* OP50 (Lewis and Fleming 1995; Brenner 1974).

### Molecular biology

For *azyx-1* localisation, 3474 bp upstream of the *azyx-1* stop codon were amplified by PCR from wild-type genomic DNA, along with 558 bp of *azyx-1* 3’UTR and fused to 5’ and 3’ ends of mNeonGreen (minus its start-AUG) using HiFi DNA assembly (NEBuilder®). The resultant linear transgene was purified (Wizard® Genomic DNA purification kit, Promega) and confirmed by sequencing (oligos p001-p006, Supplemental Table. S3), and injected into wild type N2 to generate *C. elegans* strain LSC1959 (see ‘Transgenesis’ and Supplemental Table. S1). For overexpression and rescue strains (LSC1950, LSC1951, LSC1960, LSC1997, LSC1999, LSC2001; Supplemental Table. S1), a 757 bp promoter region upstream of *azyx-1* was amplified, as were 535 bp of its 3’UTR. Next, these were fused to 5’ and 3’ ends of a 587 bp synthesized *azyx-1* gBlock™ (Integrated DNA Technologies (IDT); containing all *azyx-1* exons and its first intron, with the ATG at the *zyx-1a* start mutated to CTG) using HiFi DNA assembly (NEBuilder®) and confirmed by sequencing (oligos p007-p010, Supplemental Table. S3). To build the neuronal marker transgene, mCherry was fused to 1800 bp of the *unc-47* promoter region and 497 bp of the *unc-47* 3’UTR using HiFi DNA assembly (NEBuilder®) and confirmed by sequencing (oligos p011-p016, Supplemental Table. S3).

### CRISPR/Cas9-mediated knockout of *azyx-1a* (*lst1687*)

For the generation of the *lst1687* allele, which contains a 27 bp deletion at the beginning of the *azyx-1* ORF, the *dpy-10* co-CRISPR strategy was used with homology-directed repair (HDR) according to (Paix et al. 2017). The injection mix comprised 2.5 μl of recombinant codon-optimized Cas9 enzyme, 2.5 μl tracrRNA (0.17 mol/l, IDT), 1 μl *dpy-10* crRNA (0.6 nmol/μl, IDT), 1 μl of *azyx-1* crRNA (0.6 nmol/μl, IDT, Supplemental Table. S3), 1 μl *dpy-10* repair template (0.5 mg/ml, Merck) and 1 μl repair template for *azyx-1* containing a 27 bp deletion that encompasses *azyx-1a* start codon (1 mg/ml, IDT, Supplemental Table. S3). The mix was micro-injected in the gonads of young adult N2 Bristol wild types (Zeiss Axio Observer A1 with Eppendorf Femtojet and Eppendorf Injectman NI2)(Mello et al. 1991). Offspring were screened for the desired CRISPR/Cas9-mediated deletion by SspI (FastDigest™ Thermo Fisher) cleavage of PCR products of the *azyx-1* locus, which is only possible after HDR (oligos p007 and p0017, Supplemental Table. S3). The presence of the homozygous *lst1687* allele was confirmed via sequencing and the *azyx-1a* deletion mutant strain LSC1898 (Supplemental Table. S1) was isolated.

### Sample collection and preparation for proteomics

Adult worms were synchronized by standard hypochlorite treatment (Porta-de-la-Riva et al. 2012). After overnight incubation in S-basal (5.85 g NaCl, 1 g K2 HPO4, 6 g KH2PO4 in 1 L milliQ) on a rotor at 20 °C, the L1 arrested animals were grown on NGM plates seeded with *E. coli* OP50. For wild-type sampling at different ages, we collected worms at larval (L4, 48h post L1 refeeding), day 1 adult (20h post L4 harvest) and post-reproductive, day 8 of adulthood stages. For day-8 samples, offspring were avoided by supplementing the worm cultures with 50 μl of a 50 μM fluorodeoxyuridine (FUDR) solution every 48h, as of the L4 larval stage (*i*.*e*. L4, and days 2, 4 and 6 of adulthood) (Mitchell et al. 1979). For comparisons of wild types with *azyx-1* deletion mutants, both strains were synchronized and then grown until the day-1 adult stage. For sampling, worms were washed off NGM plates with S-basal and allowed to settle in conical tubes for 10 min. Following that, the supernatant was removed and worm pellets were diluted to 15 ml in S-basal for sorting. Worms were sorted using a Complex Object Parametric Analyzer and Sorter (COPAS) platform (Union Biometrica, Holliston, MA, USA) for each sample individually. Four independently grown populations of worms were used per condition. We collected 200 animals per sample for day-8, or 1000 animals per sample for all other conditions into a 1.5 ml Eppendorf LoBind™ tube. The worms were pelleted by spinning at 1,500 *g* for 1 min, S-basal was removed, and 200 μl of 50 mM HEPES were added to the worm pellet, spun at 1,500 *g* for 1 min and the supernatant was carefully discarded ensuring no worms were lost in pipetting. Finally, the pellet was supplemented with 1 fmol/worm of synthetic spike-in peptide (EAVSEILETSRVSGWRLFKKIS), comprising a proteotypic peptide for quantitation (EAVSEILETSR) (Vandemoortele et al. 2016) fused to a HiBit Tag (VSGWRLFKKIS) via a tryptic cleavage site, from a stock solution in water at a concentration of 100 fmol/μl. The pellet was snap frozen in liquid nitrogen and stored at -80 °C until further processing. The duration from initial worm collection off NGM until snap freezing was approximately 20 mins and carried out at 20 °C.

For protein extraction, worm pellets were thawed on ice with 100 μl of lysis buffer (8 M Urea, 2 M Thiourea in 10 mM HEPES) and lysed by sonication using a probe sonicator (40% amplitude, 5 sec ON, 10 sec OFF x 10). The lysate was spun at 15,000*g* for 10 min and the supernatant was transferred to a fresh 1.5 ml Eppendorf LoBind™ tube. Protein concentration was estimated using a Bradford assay and sample aliquots corresponding to 50 μg of total protein were processed further for LC-MS/MS. For this, each sample was reduced with 5 mM dithiothreitol at 56 °C for 30 min and alkylated with 25 mM of iodoacetamide for 20 min at room temperature. The lysate was digested overnight at 37 °C with 2 μg of sequencing-grade trypsin (Promega), after which the sample was acidified to 0.1% formic acid, cleaned using Pierce™ C18 spin columns as per the manufacturer’s protocol and dried in a Savant SpeedVac. The dried peptides were dissolved to 0.1 μg/μl in 2% acetonitrile/98% H2O/0.1% formic acid (FA)/0.1X Biognosys iRT peptides (for retention time calibration).

### Peptide ion selection for targeted quantification

Peptide ions useful for quantification of proteins of interest (ZYX-1 and AZYX-1), and of proteins used for data normalisation (GPD-3, HIS-24, spike-in) were selected based on an unscheduled parallel reaction monitoring (PRM) experiment. To accurately normalize data across age and conditions, we chose 3 normalization options: two relying on endogenous *C. elegans* proteins - *viz*. GPD-3 (GAPDH homolog - 4 peptides), HIS-24 (Histone homolog - 4 peptides) - and one relying on the externally added synthetic spike-in peptide (1 peptide). Skyline-daily was used to build an initial library (Pino et al. 2020). For all proteins of interest, all theoretically predicted tryptic peptides with a length between 7 and 26 amino acids were added to the initial spectral library. In total, 98 peptide precursor ions were selected and measured in an unscheduled PRM experiment, that was run on a pooled sample consisting of all peptide samples used in this study and analysed with Skyline. Subsequently, 23 measured peptide precursors representing 22 peptides and 5 target proteins (ZYX-1, AZYX-1, GPD-3, HIS-24, Spike-in) were selected for the final PRM measurements. Additionally, 11 MS1 ions of the Biognosys iRT reference peptides were included in the precursor list. Details of all peptides and corresponding protein(s), including their uniqueness in the proteome database, can be found in Supplemental Table. S2.

### Targeted LC-MS/MS measurements

Targeted measurements using scheduled PRM were performed with a 50 min linear gradient on a Dionex UltiMate 3000 RSLCnano system coupled to a Q-Exactive HF-X mass spectrometer (Thermo Fisher Scientific). The spectrometer was operated in PRM and positive ionization mode. MS1 spectra (360–1300 m/z) were recorded at a resolution of 60,000 using an AGC target value of 3×10^6^ and a MaxIT of 100 ms. Targeted MS2 spectra were acquired at 60,000 resolution with a fixed first mass of 100 m/z, after HCD with a normalised collision energy of 26%, and using an AGC target value of 1×10^6^, a MaxIT of 118 ms and an isolation window of 0.9 m/z. In a single PRM measurement, 23 + 11 MS1 peptide ions (see above) were targeted with a 5 min scheduled retention time window. The cycle time was ∼2.1 s, which leads to about 10 data points per chromatographic peak.

### Targeted mass spectrometric data analysis

PRM data were analysed using Skyline (version 64-bit 21.1.0.278) (Pino et al. 2020). Peak integration, transition interferences and integration boundaries were reviewed manually, considering 4-6 transitions per peptide. To discriminate true from false positive peptide detection, filtering according to correlation of PRM fragment ion intensities was carried out. For this purpose, an experimental spectral library was built from the PRM data itself, by searching these with MaxQuant and then loading the generated search results back into Skyline. For confident peptide identification, a “Library Dot Product” ≥0.85, as well as a mass accuracy ≤10 ppm (“Average Mass Error PPM”) were required. We also manually verified the correlation between PRM fragment ion intensties and spectra predicted with the artifical intellegence algorithm Prosit (Gessulat et al. 2019). For peptide and protein quantification, chromatographic peak areas were exported from Skyline in MSStats format, and further processing, quantification, statistical analysis and visualisation were performed in RStudio with the MSStats package (Choi et al. 2014). For HIS-24, 3 most consistent peptides out of 4 were considered for downstream analysis and peptide FISQNYK was omitted. The data were log2 transformed, processed as per default MSStats parameters and visualized using the ggplot2 package of *R*. For age analysis, data were normalized to spike-in peptide (1 fmol/worm) and L4 samples were used as the reference. For *azyx-1a* mutant and wild type comparison, all three normalizations were considered (*i*.*e*., GPD-3, HIS-24 and spike-in). The mass spectrometric raw files acquired in PRM mode and the Skyline analysis file have been deposited to Panorama Public (Sharma et al. 2018) and can be accessed via https://panoramaweb.org/Peu1.url.

### Transgenesis

For *in vivo* localisation of *azyx-1*, the *lstEx1065* extra-chromosomal array was generated by mixing 25 ng/μl purified linear transgene [*azyx-1p::azyx-1*+mNeonGreen::*azyx-1* 3’UTR] (see ‘Molecular biology’) with 25 ng/μl coelomocyte-restricted co-injection marker [*unc-122p::DsRed*] and 50 ng/μl 1-kb ladder (Thermo Scientific) as carrier DNA. This was injected into Bristol wild type (N2) to generate LSC1959 (Supplemental Table. S1).

All genetic overexpressions and rescues of *azyx-1* strains (Supplemental Table. S1) were created via injection of 25 ng/μl of linear transgene [*azyx-1p::azyx-1*(gBlock)::*azyx-1* 3’UTR] with 12.5 ng/μl pharyngeal co-injection marker [*myo-2p*::mCherry] and 50 ng/μl of 1-kb ladder (Thermo Scientific) as carrier DNA. For strains transgenically expressing the *unc-47p*::mCherry neuronal marker (LSC1998 and LSC1999; Supplemental Table. S1), 10 ng/μl of this linear construct was injected along with 5 ng/μl pharyngeal co-injection marker [*myo-2p*::mCherry] and 50 ng/μl of 1-kb ladder (Thermo Scientific), with the addition of 10 ng/μl overexpression transgene [*azyx-1p*::*azyx-1*(gBlock):: *azyx-1* 3’UTR] for LSC1999.

Transgenic strains were always confirmed by observation of co-injection marker presence via fluoresence microscopy, followed by PCR and sequencing of the added target sequences. For overexpression of *azyx-1* in the *zyx-1* reporter background (LSC1870, which expresses mCherry::*zyx-1*;*zyx-1*::GFP, Supplemental Table. S1), the extrachromosomal array (NM3425) was integrated with UV irradiation as per (Mariol et al. 2013) and outcrossed twice with wild type (N2 Bristol). All injections using a Zeiss Axio Observer A1 with Eppendorf Femtojet and Eppendorf Injectman NI2 were performed targeting syncytial gonads of young adults.

### Confocal imaging

Confocal microscopy was performed using either an Olympus FluoView 1000 (IX81) or a Zeiss LSM900 confocal microscope. To obtain synchronized L4 larvae, a timed egg-laying was performed 48 hours before imaging whereas day 1 adults were synchronized by picking L4 larvae approximately 16 hours before imaging. Worms were anaesthetized using of 500 mM sodium azide and mounted on 2% agarose pads. Z-projections of representative images were made using Fiji (Schindelin et al. 2012).

### Manual scoring of muscular filaments

The image files for day 1 adult RW1595 (*cohort 1 = 5, cohort 2 = 24), LSC2000 (cohort 1 = 12, cohort 2 = 20) and LSC2001 (cohort 1 = 16, cohort 2 = 22)* were randomized in Blinder freeware (Cothren et al. 2018) and scored in three qualitative classes of muscle filaments: (1) Normal well-organized, (2) moderately disorganized and (3) severely disorganized (Fig. 6A) based on reported manual scoring parameters (Dhondt et al. 2021).

### Burrowing assay

Burrowing assays were performed with synchronized adults and executed essentially as described by Lesanpezeshki *et al*., (2019), with minor adjustments. Briefly, 20 gravid adults for each replicate were allowed to lay eggs on a seeded NGM plate for 3 hours, and subsequently removed while allowing the eggs to hatch and grow at 20 °C. After 70 hours, worms were washed off the NGM plates with S-basal and transferred onto unseeded NGM plates to induce a starvation response. After 1 hour, 30 adult worms were dropped in 10 μl S-basal in a well of a Corning Costar® 12-well plate and covered with 2.3 ml of 26% w/v Pluronic F-127 (Sigma-Aldrich) at 14 °C. After 15 min at 20 °C, the Pluronic F-127 had gelated and a droplet of 20 μl 100 mg/ml *E. coli* HB101 was added on top as a chemoattractant (time = 0). The bacterial droplet was monitored every 30 mins for 3 hours to calculate a chemotaxis index as the percentage of worms that had cumulatively reached the bacteria (out of the 100% corresponding to n=30). At each time point of observation, worms that had reached the bacterial pellet were removed to avoid crowding and reburrowing. Statistical significance was determined by two-way ANOVA.

## Acknowledgements

This work was supported by FWO Flanders (grants G052217N), KU Leuven (C16/19/003) and EPIC-XS, grant number 823839, funded by the Horizon 2020 programme of the European Union. Some strains were provided by the CGC, which is funded by NIH Office of Research Infrastructure Programs (P40 OD010440). We would also like to thank Prof. Kathrin Gieseler (Université Claude Bernard Lyon 1, France), Prof. Michael Nonet (Washington University in St. Louis, USA) and Prof. Bart Braeckman (UGent, Belgium) for providing strains, and Marlies Peeters and Dr Gerben Menschaert (UGent, Belgium) for valuable discussions.

